# Laminin-driven Epac/Rap1 regulation of epithelial barriers on decellularized matrix

**DOI:** 10.1101/612325

**Authors:** Bethany M. Young, Keerthana Shankar, Cindy K. Tho, Amanda R. Pellegrino, Rebecca L. Heise

## Abstract

Decellularized tissues offer a unique tool for developing regenerative biomaterials or *in vitro* platforms for the study of cell-extracellular matrix (ECM) interactions. One main challenge associated with decellularized lung tissue is that ECM components can be stripped away or altered by the detergents used to remove cellular debris. Without characterizing the composition of lung decellularized ECM (dECM) and the cellular response caused by the altered composition, it is difficult to utilize dECM for regeneration and specifically, engineering the complexities of the alveolar-capillary barrier. This study takes steps towards uncovering if dECM must be enhanced with lost ECM proteins to achieve proper epithelial barrier formation. To achieve this, epithelial barrier function was assessed on dECM coatings with and without the systematic addition of several key basement membrane proteins. After comparing barrier function on collagen, fibronectin, laminin, and dECM in varying combinations as an *in vitro* coating, the alveolar epithelium exhibited superior barrier function when dECM was supplemented with laminin as evidenced by trans-epithelial electrical resistance (TEER) and permeability assays. Increased barrier resistance with laminin addition was associated with upregulation of Claudin-18, E-cadherin, and junction adhesion molecule (JAM)-A, and stabilization of zonula occludens (ZO)-1 at junction complexes. The Epac/Rap1 pathway was observed to play a role in the ECM-mediated barrier function determined by protein expression and Epac inhibition. These findings reveal potential ECM coatings and molecular therapeutic targets for improved regeneration with decellularized scaffolds or edema related pathologies.

## 1. INTRODUCTION

Decellularized lung tissue engineering offers a promising regeneration strategy for patients with a variety of incurable pulmonary diseases. This technology can provide the native structure and biochemical cues of the lung to guide healthy maturation of functional tissue. Decellularized lungs are the gold standard scaffold for whole lung bioengineering; however, current approaches do not produce a long-term, functional lung replacement, indicated by severe edema after implantation. Insufficient recellularization and immature barriers keep this technology from the clinic. Most of the research to date has focused on assessments of stem cell differentiation and attachment [1–4] or mimicking lung development during regeneration [5,6]. More recently, researchers in this field have examined recellularization on a smaller scale by evaluating endothelial barriers [7]; however, there is little research on epithelial barrier formation with recellularized lungs. This research takes a unique approach to re-epithelialization of whole organ scaffolds by examining alveolar junction formation on a cellular level to then inform future recellularization strategies. Therefore, we are attempting to systematically determine which components are essential to alveolar barrier function and develop a scaffold coating to replenish these key matrix proteins.

Homeostatic alveolar epithelial barrier function is maintained by two main structures: tight junctions (TJ) and adherens junctions (AJ). Both junctions provide tension resistance and selective permeability during resting mechanical load. Formation of strong barriers relies on a highly regulated sequential recruitment of several main junction proteins including cadherins, claudins, zonula occludins (ZO), and junctional adhesion molecules (JAM)s [8–10]. Protein kinase A (PKA) and exchange protein directly activated by cAMP (Epac) have both been identified as regulators of barrier function by activation of G-protein-coupled receptor and release of cyclic AMP (cAMP) [11–15]. While PKA has traditionally been studied as the primary mediator of barrier function, inhibition of PKA does not result in substantial barrier disruption [16–18]. This has led to a recent transition into investigating the role of Epac within endothelial [18–23] and less often, epithelial barriers [24–26]. After activation, Epac is rapidly translocated to the plasma membrane from the cytoplasm and nuclear membrane to activate Rap1 to regulate afadin (AF-6) stabilization of AJs, TJs, and integrins [27]. Within epithelial cells, Epac signaling has also been implicated in inhibition of cell migration and epithelial to mesenchymal transition (EMT), [28] and promoting adhesion to laminin and fibronectin [29–31]. This suggests that epithelial junction mediation may be a function of Epac in response to specific ECM proteins; however, the direct causes and mechanisms of this within the alveolar epithelium are not fully understood.

Decellularization techniques utilize several harsh detergents to ensure complete removal of cellular debris, but these detergents can drastically alter the ratio of ECM components that are left behind. Matrix proteins such as collagens, elastin, laminin, fibronectin, and glycosaminoglycans (GAGs) are preserved through the decellularization process, but there are reports of up to a 50% loss of most of these components compared to native ECM [32,33]. Variations in ECM composition and stiffness can have a profound effect on cell phenotype, as well as cell attachment and barrier function [34–36]. High collagen and fibronectin content promote cell attachment at the expense of suboptimal barriers, indicative of metastasis and fibrosis [37–40]. Laminin is integral to both strong barrier formation and epithelial differentiation [41–47]. We hypothesize that the discrepancy between native ECM and decellularized airway surface ECM can alter the epithelial cell-cell junction assembly during recellularization through activation of Epac. Understanding how and which ECM components modulate alveolar barriers, specifically TJs and AJs, is integral to producing a whole lung replacement that is functional on the cellular level. This research is the first step to creating a tailorable ECM environment within decellularized lungs by methodically replenishing basement membrane proteins that promote alveolar barrier formation.

## 2. RESULTS

### 2.1 Production and characterization of lung dECM to investigate epithelial barrier function

Lung dECM coatings were produced with porcine lungs decellularized over 3 days with Triton X-100, sodium deoxycholate (SDC), and DNAse as described previously [48]. The acellular tissue was lyophilized and milled into a powder that could be solubilized by pepsin in 0.01M HCl for 4 h. Coatings were supplemented with either laminin or fibronectin before culturing mouse alveolar epithelial cells (MLE12s) or basal epithelial stem cells (BESCs) to assess barrier function outcomes (Fig. 1A). ECM produced by decellularization retains many of the critical matrix proteins within the lung; however, it is unknown whether any of these ECM components have been damaged or partially removed by the decellularization process. We then sought to determine the effect of dECM on epithelial response compared to cells cultured on laminin or fibronectin alone or in combination with dECM.

**Fig. 1.**
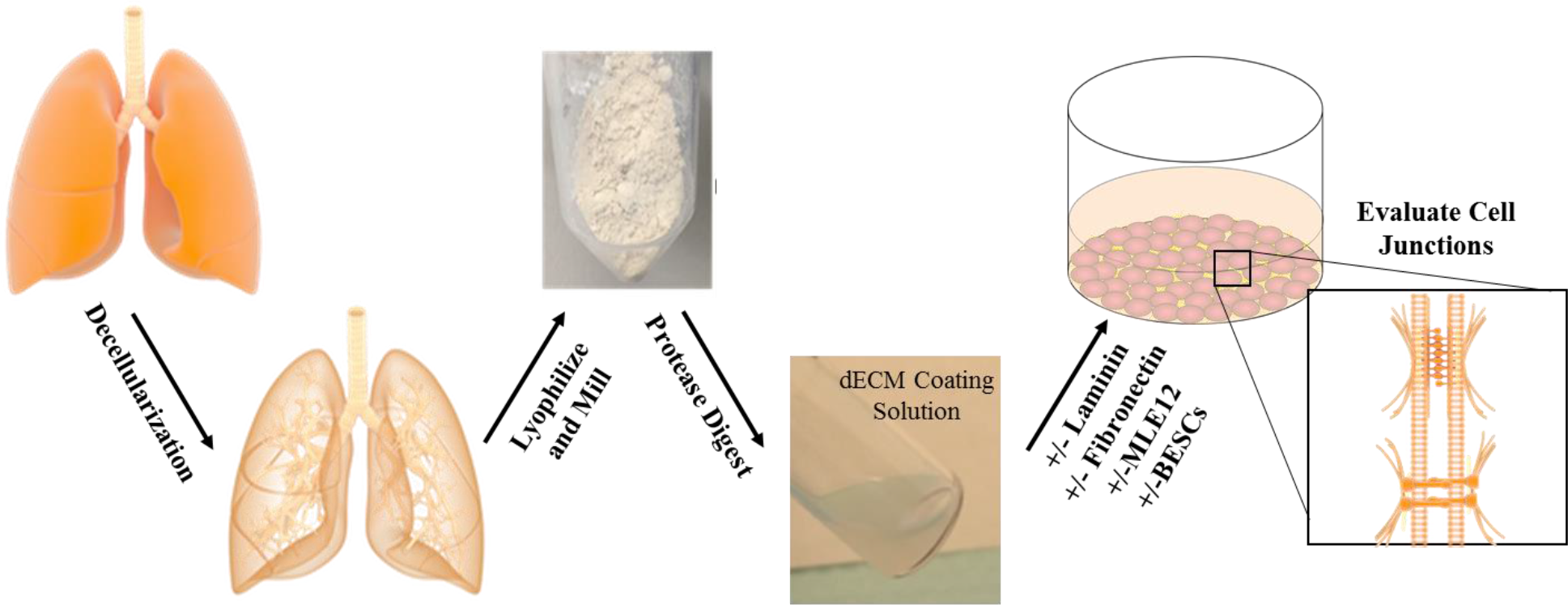
Experimental design of dECM production, coating procedures, and composition. Porcine lungs are decellularized and milled into a powder that can be solubilized and used as a coating solution. The coating is supplemented with barrier-mediating ECM proteins to evaluate TJ and AJ of the alveolar barrier formed by MLE12 cells and BESCs.

### 2.2 Decellularized lung ECM requires supplementation for proper epithelial adhesion and barrier formation

Alveolar epithelial cell barrier formation on each dECM combination compared to dECM alone was examined by TEER and transwell permeability of MLE12 cells cultured onto matrix-coated plates (Fig. 2A-B). The first 12 h indicates resistance predominately caused by attachment and spreading of the cells on each matrix, showing initially increased resistance by cells cultured on laminin alone, dECM with laminin, and dECM with fibronectin coatings. Cell viability at 4 and 24 h using CCK8 also shows a significant increase of MLE12 attachment to fibronectin-enhanced matrices compared to all other substrates (Supplemental Fig.1). This trend persisted through 24 h when the highest resistance monolayer was produced by cells cultured on the combination of dECM and laminin. dECM with laminin has a significantly higher barrier function compared to collagen and fibronectin for the entire 3-day culture.

**Fig. 2.**
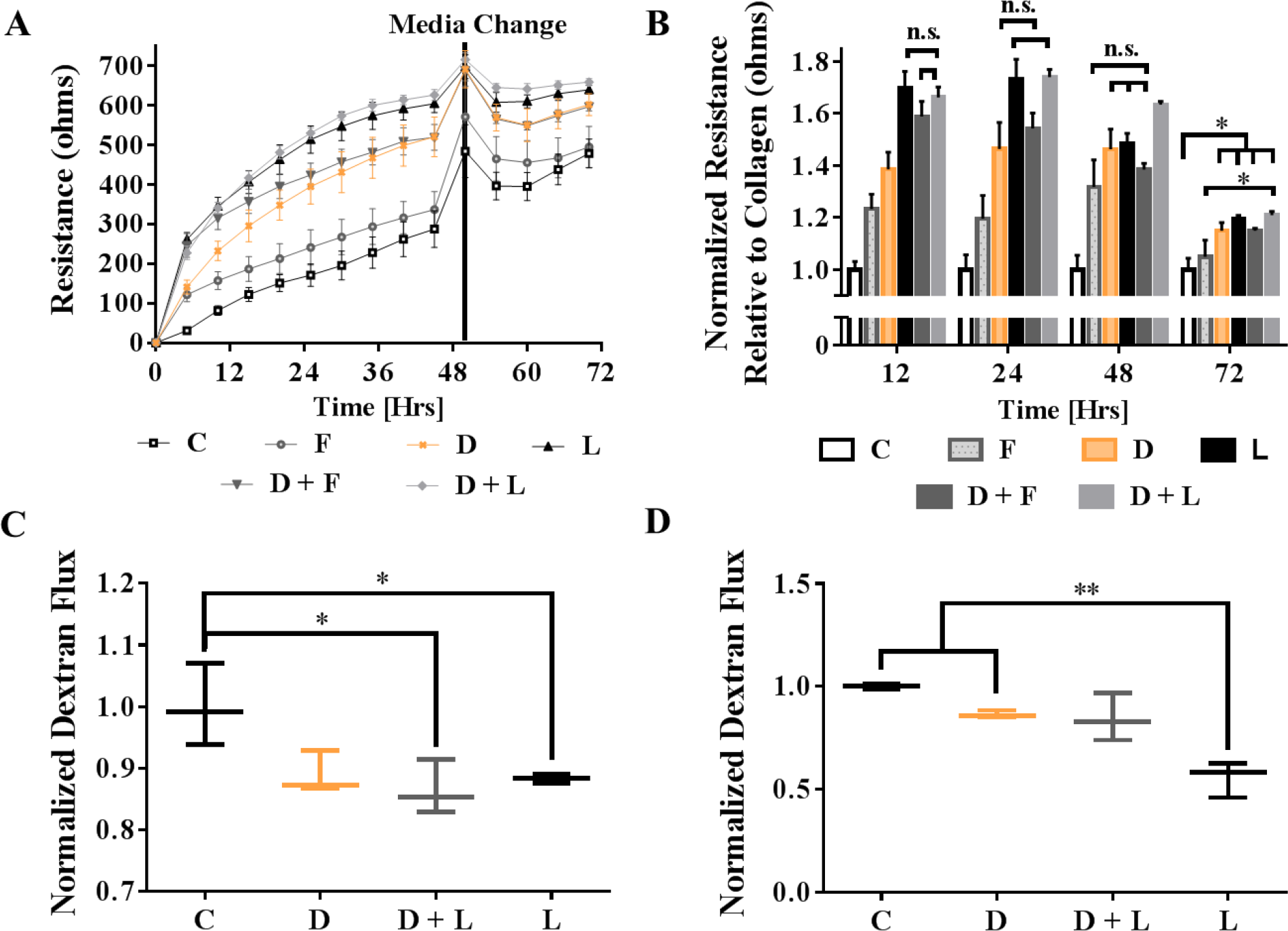
The effect of dECM composition on alveolar epithelial barrier formation. TEER of MLE12 alveolar epithelial cells cultured on TCP coated with collagen (C), dECM (D), laminin (L), fibronectin (F) or a combination of two for 72 h is shown by (A) a representative real-time graph of one sample with 3 experimental replicates and (B) binned at 12, 24, 48 and 72 h with 3 experimental replicates to show statistical differences. MLE12 monolayer permeability to small molecules with respect to ECM coating after (C) 2 and (D) 4 days of cultures was determined by a 4 kDa FITC-Dextran Transwell permeability assay. Normalized dextran flux is shown as absorbance values from the bottom chamber of the Transwell, correlating to dextran transported across the membrane. Data are normalized to the collagen control and presented as mean +/− standard deviation. N ≥ 3 for each coating condition. n.s, *, and ** indicates p > 0.05, p < 0.05, and p < 0.001, respectively.

Small molecule permeability with 4 kDa dextran by MLE12 monolayers formed onto matrix-coated transwell supports was also examined. At 2 days of culture, both dECM + laminin and laminin alone have less permeability compared to collagen, but not significantly different from dECM (Fig. 2C). It is only after four days that cells cultured on laminin alone are less permeable compared to cells cultured on both collagen and dECM (Fig. 2D).

### 2.3 Analysis of ECM-mediated AJ and TJ protein recruitment within alveolar epithelium

Increased barrier function with laminin led us to investigate the underlying mechanisms further. MLE12s were seeded on ECM coatings for 5 days to evaluate mRNA expression with qPCR (Fig. 3A-F). The laminin concentration of 0.1 mg/mL was chosen based on decreased junction gene expression with higher coating concentrations (data not shown). We concluded that 0.1 mg/mL laminin is the most effective concentration for further experimentation. Gene expression of Claudin 18, Epac, and AF-6 all increased with cells cultured onto laminin-supplemented matrices. E-cadherin and JAM-A gene expression were highest with the combination of laminin and dECM. Rap1 expression only had significant differences between cells cultured on dECM with laminin and dECM with fibronectin. More significant differences between other coating groups may be seen in Rap1 localization or activation. Expression of all genes were either decreased or not significantly different by cells cultured on coatings containing laminin compared to all other coating groups. Additionally, there is an increasing trend in expression of AF-6, JAM-A, and Rap1 with cells cultured on dECM compared to on collagen alone, suggesting that collagen alone is not enough to stimulate this pathway and the remnants of laminin or other proteins trigger Epac/Rap1 activation.

**Fig 3.**
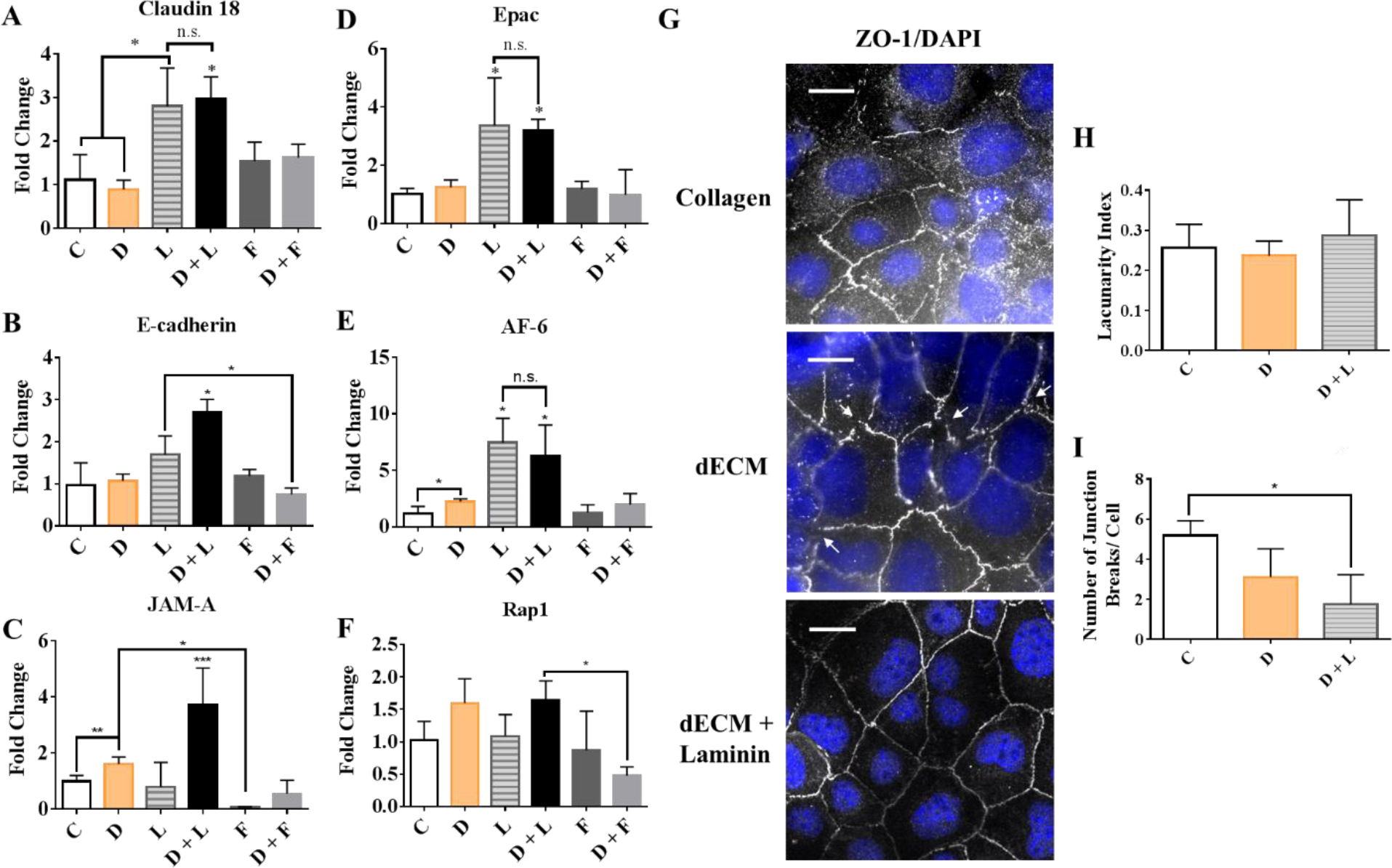
Changes in junctional gene expression and ZO-1 localization with dECM coatings dictates alveolar epithelial junction regulation. MLE12 alveolar epithelial cells were cultured on TCP coated with collagen (C), dECM (D), laminin (L), fibronectin (F) or a combination of two for 5 days before collecting mRNA. Gene expression of the (A-C) junction proteins, claudin-18, E-cadherin, and JAM-A, and (D-F) junction regulators, Epac, AF-6, and Rap1 were quantified using qPCR and normalized to collagen coating controls. Upon barrier formation by alveolar epithelial cells, (G, scale bar = 20 μm) immunofluorescent staining of ZO-1 (white) and Dapi (blue) shows stabilization of the junctions depending on coating type by a representative confocal image. Arrows indicate breaks within the cell junctions. Quantification of confocal images from 3 experimental replicates using AngioTool determined the (H) lacunarity index and (I) the number of junction interruptions per cell. Data are presented as mean +/− standard deviation. N ≥ 3 experiments with 2-3 technical replicates for each coating condition. n.s, *, and ** indicate p > 0.05, p < 0.05, and p < 0.001, respectively.

ZO-1 localization by immunofluorescence staining (Fig. 2G) shows a qualitative increase in the amount of ZO-1 stabilizing the junctions of MLE12s cultured on dECM with laminin and laminin alone compared to collagen. Compared to cells cultured on collagen, there is also more uniformity in cell and junctional morphology of MLE12s on both dECM and dECM with laminin coatings. Junctions formed on dECM or collagen coatings both exhibit gaps within the ZO-1 lining the junction, (shown by arrows,Fig. 3G), and a zipper arrangement of the ZO-1 at the junction. Alternatively, laminin coatings appear to induce a more continuous, linear ZO-1 formation at the junction Quantitative analysis by ImageJ shows no significant difference in the lacunarity index (Fig. 3H), corresponding to cell junction shape uniformity, and a decrease in the number of junctions per cell with laminin added to the coating before culture.

### 2.4 Epithelial progenitor cell barrier formation

To determine if similar findings would translate to a progenitor cell population that is commonly used for lung tissue engineering, basal epithelial stem cell (BESC) junction formation through the Epac/Rap1/AF-6 pathway was probed. BESCs were cultured onto collagen, dECM, and dECM with laminin-coated TCP for a week prior to examination with TEER (Fig. 4A-B). Initially, there were few differences in resistance means between BESCs cultured on dECM and dECM with laminin coatings. Changes were observed after 168 h when the barrier function of cells cultured on dECM + Laminin begin to reach higher resistances overall. Collagen shows very low resistance throughout the entire culture, caused by lack of BESC junction formation and persistence of a round morphology.

**Fig 4.**
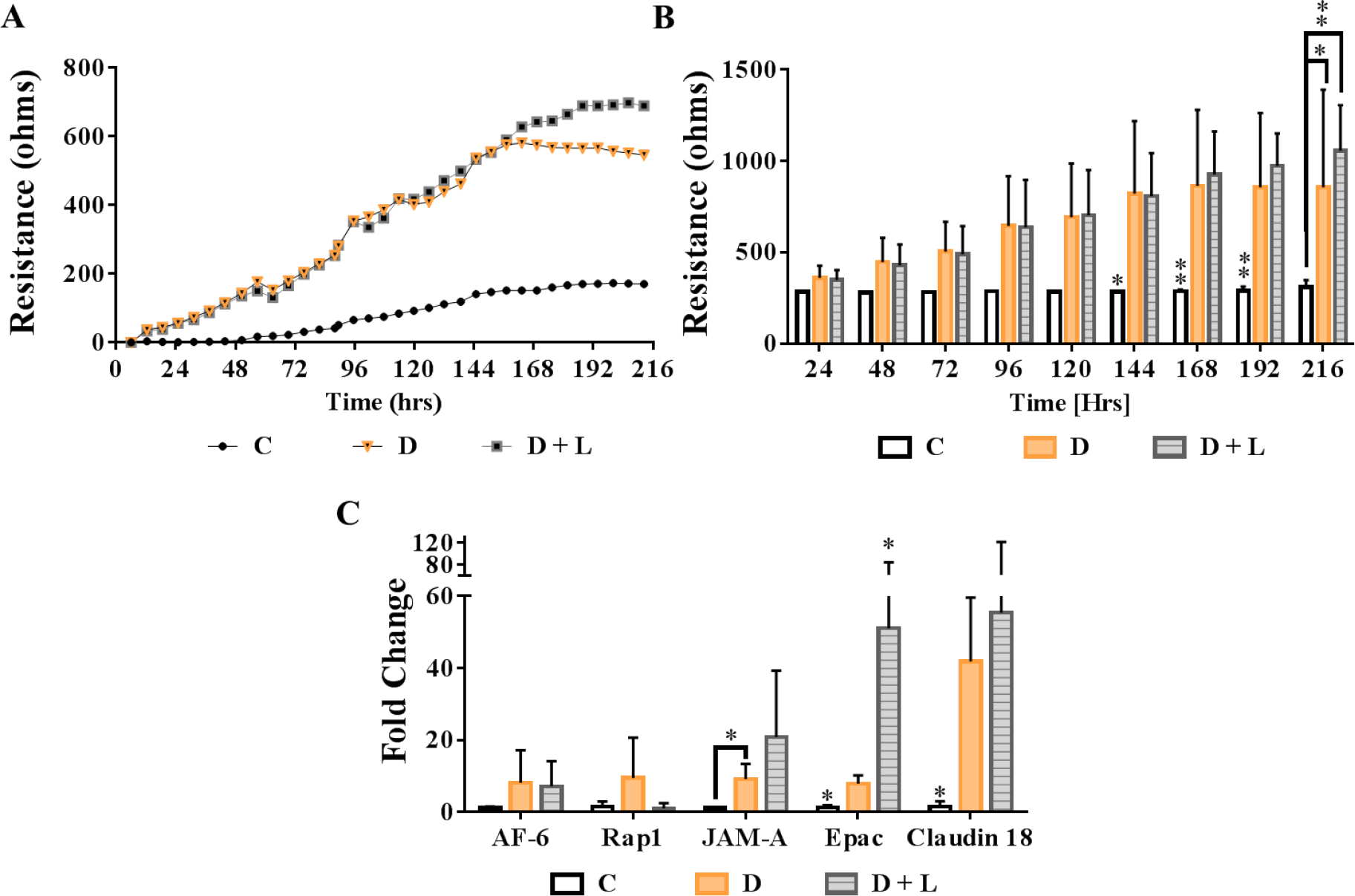
BESC barrier function and Epac/Rap1 pathway gene expression. TEER of BESCs cultured on TCP coated with collagen (C), dECM (D), laminin (L) or a combination of two was examined over 9 days after being cultured on collagen, dECM or dECM + Laminin coatings for 1 week. (A) A representative graph with adjusted means and (B) binned into 24-hour periods with raw resistances is shown. (C) Gene expression of AF-6, Rap1, JAM-A, Epac, and Claudin- 18 after 5 days of culture on each respective coating quantified with qPCR. Unless otherwise stated, all data are presented as mean +/− standard deviation. N ≥ 3 for each coating condition. * and ** indicates p < 0.05, and p < 0.001, respectively.

Similar to MLE12s cultured on laminin coatings, dECM causes significant upregulation of JAM-A and Claudin-18, but not in Epac. Contrasting MLE12 gene expression, BESCs do not show significant increases in AF-6 expression when cells are cultured on dECM and dECM with laminin compared to collagen.

### 2.5 Role of Epac in laminin-mediated barrier formation

To fully understand the effects of the Epac/Rap1 upregulation on laminin-rich matrices, inhibition or activation of Epac with ESI-09 or 007-AM, respectively, was used to evaluate if laminin-mediated barrier reinforcement could be abolished. TEER analysis of MLE12 cells treated with EPAC inhibitor or agonist for 24 h shows significantly decreased barrier resistance with ESI09 Epac inhibition and barrier strengthening with 007AM Epac activation over 24 h (Fig. 5A). Epac treatments were reversible upon media change, shown by the convergence of both groups with their DMSO counterpart.

**Fig. 5.**
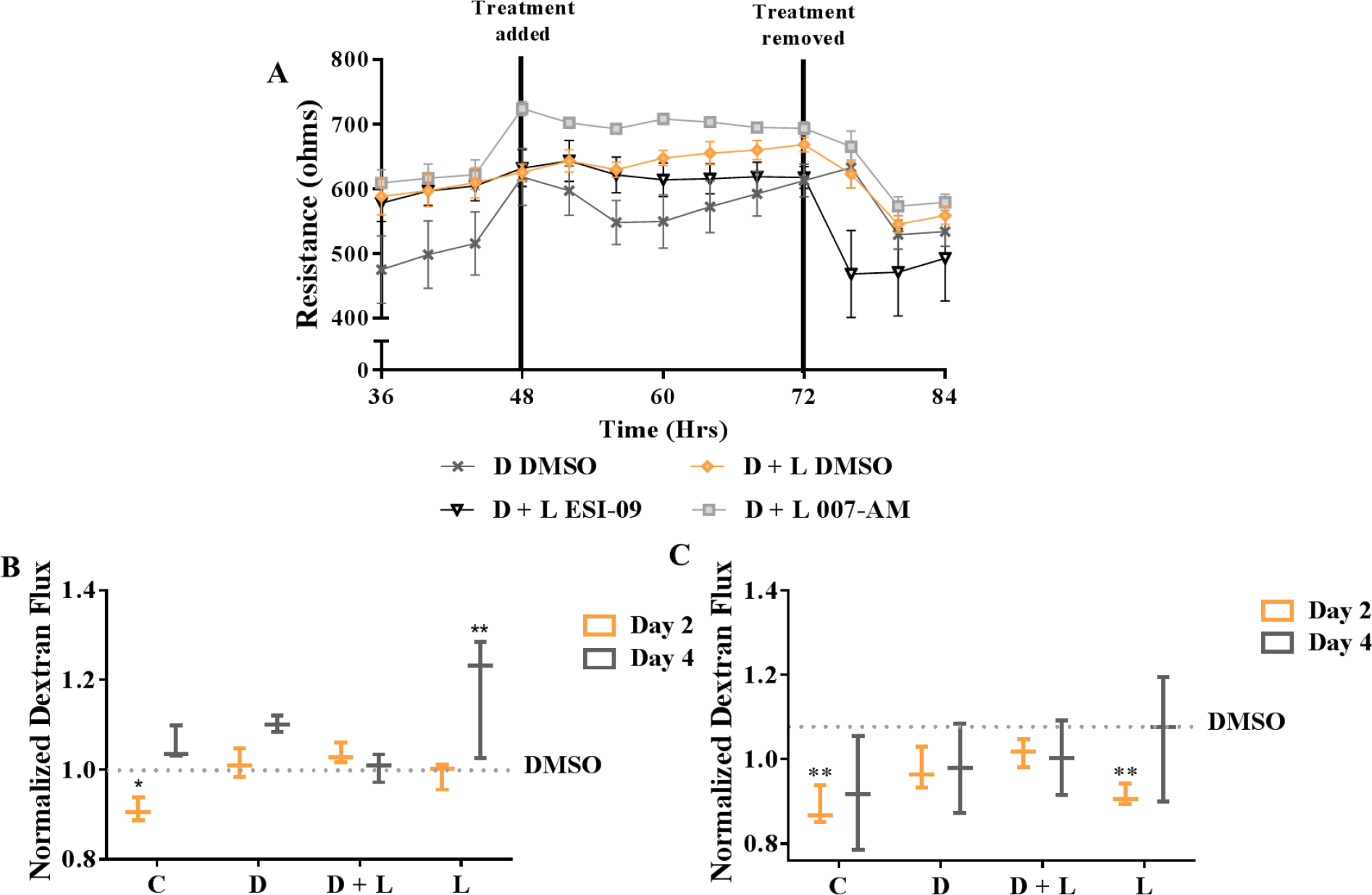
Both Epac activation and inhibition cause ECM-dependent barrier function changes. (A) MLE12 alveolar epithelial cells were cultured on TCP coated with collagen (C), dECM (D), laminin (L), or a combination of two and treated with either ESI-09 or 007-AM for 24 hours to examine TEER. (B) Cells cultured on dECM with laminin and laminin only coatings were treated with ESI-09 to determine if barrier permeability to 4 kDa molecules can be abolished. (C) 4 kDa FITC-dextran permeability of cells cultured on collagen and dECM with 007-AM treatment was evaluated to determine whether barrier permeability of 4 kDa molecules could be further reduced. All data are presented as mean +/− standard deviation. N ≥ 3 for each coating condition. * and ** indicates p < 0.05, and p < 0.001, respectively.

While TEER of cells cultured on dECM + Laminin can be reduced with ESI-09, 4 kDa dextran permeability does not change significantly from DMSO (Fig. 5B). However, barrier disruption was observed when ESI-09 is added to MLE12 culture on laminin alone after 4 days, shown by an increase in dextran flux (Fig. 5B). An increase in barrier resistance was also quantified when ESI-09 was added to cells cultured on collagen. The reason for this is unknown, but we speculate that it could be caused by a feedback loop when laminin is not present. To determine if laminin-mediated barrier function can be induced in cells cultured without laminin supplemented coatings with Epac activation, similar dextran permeability assays were also conducted on collagen and dECM with 007-AM (Fig. 5C). Treatment of cells on collagen and laminin with 007-AM after 2 days of culture were the only groups to show significant decreases in junction permeability.

Delving further into the mechanisms of Epac we also examined junctional gene and protein expression with Epac inhibition. All junctional gene expression was decreased significantly compared to the vehicle control except for Rap1, which showed a decreasing trend when treated with ESI-09 (Fig. 6A). ZO-1 localization to the junction was also disrupted with the treatment of ESI-09 (Fig. 6B). Additionally, F-actin arrangement and cell morphology were drastically altered with increased stress fiber formation and junctional degradation. This was confirmed quantitatively by image analysis (Fig. 6C-D), showing decreased mean lacunarity with ESI-09; however, a non-significant decrease in the number of junction endpoints per cell was potentially caused by lack of junctions.

**Fig. 6.**
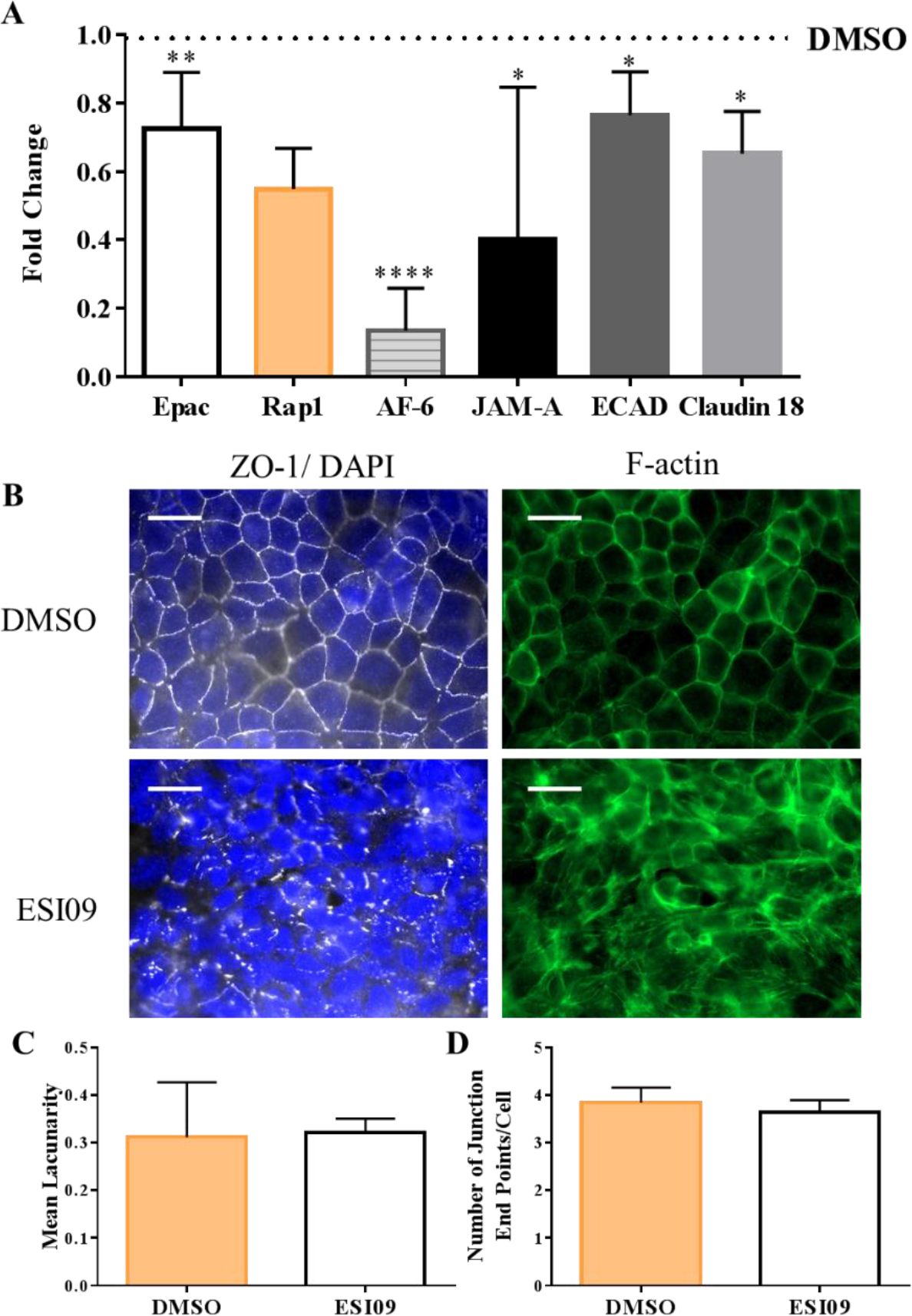
Effect of Epac inhibition on junction protein and gene expression. (A) Gene expression of claudin-18, E-cadherin, JAM-A, Epac, AF-6, and Rap1 by MLE12 alveolar epithelial cells cultured onto TCP coated with dECM with laminin for 5 days before treatment with ESI-09 was quantified and normalized to the vehicle control. (B, scale bar = 20 μm) Upon confluency, MLE12s were treated for 4 h with ESI-09 and stained for ZO-1 (white) and F-actin (green). Representative confocal images show disorganized junctions with Epac inhibition. Quantification of confocal images from 3 experimental replicates using AngioTool determined (C) the lacunarity index and (D) the number of junction interruptions per cell with ESI-09 treatment. Data is presented as mean +/− standard deviation from 3 experiments with a minimum of 2 technical replicates each and *, **, **** indicates p < 0.05, p< 0.01, and p < 0.0001, respectively.

Conversely, Epac activation with 007-AM increases gene expression of both Rap1 and JAM-A (Fig. 7 A). Immunofluorescence staining cells cultured on collagen matrices with and without 007-AM staining shows ZO-1 localization to the junctions and f-actin cortical organization (Fig. 6 B), more resembling junctions formed on laminin-rich matrices. Image quantification characterizes these changes by finding an increase in cell shape uniformity (lacunarity, Fig. 7 C) and a decrease in the number of junction-breaks per cell (Fig. 7 D). These data show that JAM-A increases expression while other AJ and TJ maturation may be caused by stabilization of ZO-1 or translocation to the junction.

**Fig. 7.**
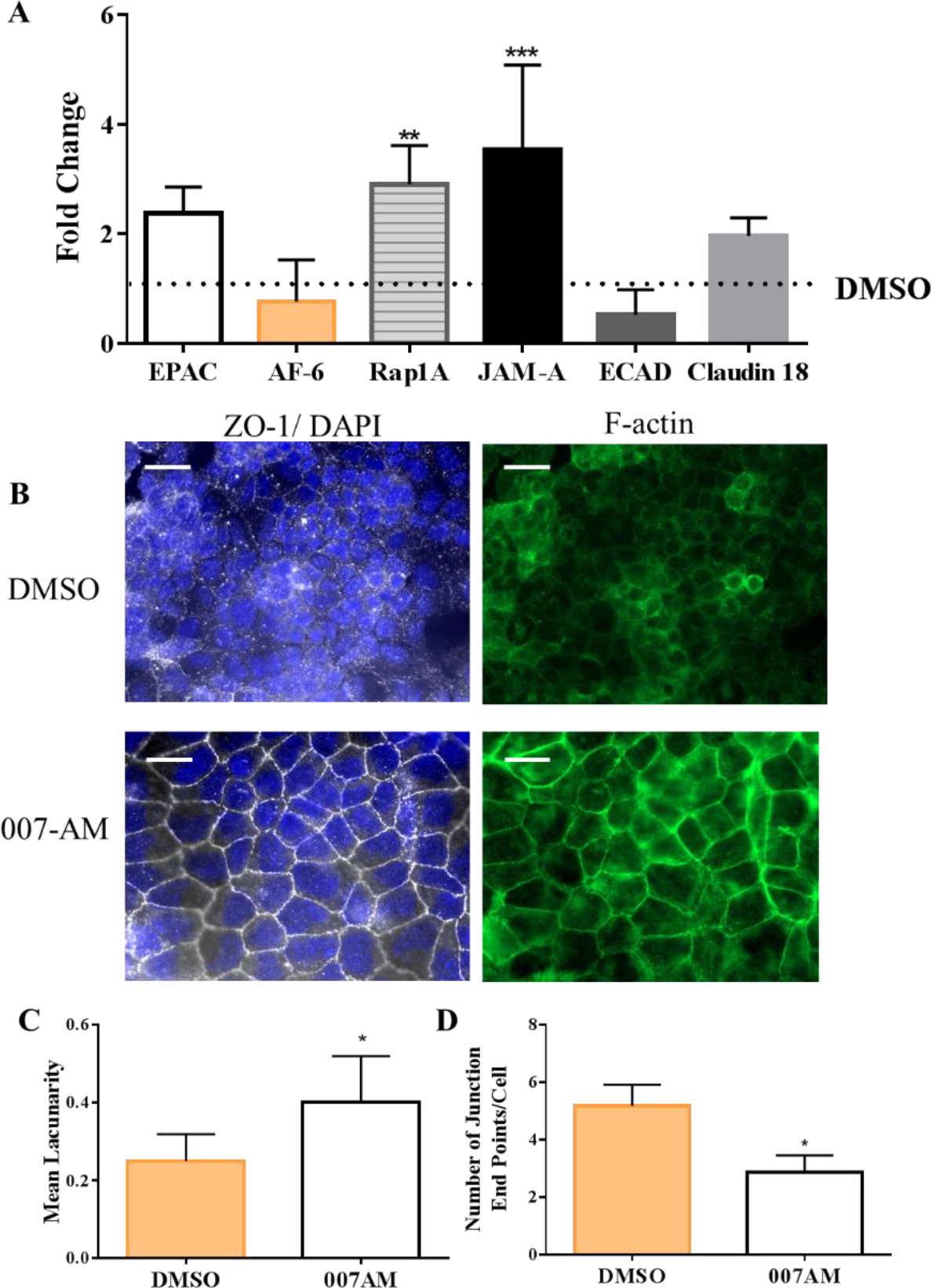
Junction reorganization with Epac agonist. (A) Gene expression of claudin-18, E-cadherin, JAM-A, Epac, AF-6, and Rap1 by MLE12 alveolar epithelial cells cultured onto TCP coated with collagen for 5 days before treatment with 007-AM was quantified and normalized to the vehicle control. (B, scale bar = 20 μm) Upon confluency, MLE12s were treated for 4 h with 007-AM and stained for ZO-1 (white) and F-actin (green). Representative confocal images show more organized junctions with 007-AM treatment. Quantification of confocal images from 3 experimental replicates using AngioTool determined (C) the lacunarity index or the uniformity of junction formation and (D) the number of junction interruptions per cell with 007-AM treatment. Data is presented as mean +/− standard deviation from 3 experiments with a minimum of 2 technical replicates each and *, **, *** indicates p < 0.05, p< 0.01, and p < 0.001, respectively.

## 3. DISCUSSION

This research aims to expand the basic knowledge of epithelial barrier physiology and to elucidate how decellularization can alter engineered alveolar barriers. To achieve this, we have investigated whether epithelial barrier function can be improved by systematically replenishing laminin or fibronectin within dECM. Laminin and fibronectin were chosen for their abundance within epithelial basement membranes and for the drastic effects both can have on epithelial differentiation and propagation of disease pathologies if concentrations are dysregulated [43,49–53].

Through TEER and dextran permeability studies, we have identified laminin as a key protein in alveolar junction formation over collagen 1 and fibronectin. Fibronectin causes a prolonged decrease in barrier function but an increase in initial attachment rate, seen both in TEER before 12 hr and CCK8 viability assays at 4 and 24 h. These results are constant with previous studies showing increased cell attachment onto fibronectin and laminin-rich matrices, but only laminin enhances barrier maturation within decellularized lungs and *in vitro* [34,54]. Analysis of epithelial barrier formation on dECM shows some similarities in the characteristics of barriers formed on both fibronectin and laminin, yet at a fraction of the resistance. At earlier time points, dECM had a similar attachment rate to fibronectin, but a long-term slope resembled cells on laminin coatings. This confirms that previously identified concentrations of both laminin and fibronectin within dECM [55] are promoting initial attachment and steady increases in barrier resistance by alveolar epithelial cells. Nevertheless, to achieve prolonged barrier maturation, laminin would need to be added.

Similarly, small molecule permeability with laminin coatings also decreased; however, dECM and dECM with laminin did not show significant changes from that of collagen, mainly after 4 days of culture. Dextran permeability showed unexpected differences from the TEER results, but we postulate that after 4 days, collagen and dECM may have formed tight enough barriers concerning pore size, although the changes in TEER may give more insight into prolonged ion movement associated with edema [56].

To further understand how laminin is increasing the barrier function of alveolar epithelium, we sought to identify which junction structures are being regulated. Claudin-18 and E-cadherin are upregulated with laminin or dECM + laminin matrices, suggesting both TJ, AJ, and focal adhesion complexes are potentially altered to achieve the overall barrier strengthening with laminin coated substrate. We believe this is due in part by the stabilization of multiple junction structures by the scaffolding protein ZO-1, seen by translocation of ZO-1 to the junction in a continuous band with laminin enriched coatings. Analysis of ZO-1 immunofluorescent staining at the cell junction showed that there was a significant decrease in the number of breaks in the ZO-1 junction staining per cell and a more linear junction morphology when cells were cultured on laminin. ZO-1 recruitment to the cell junction in an organized morphology is a strong correlation to alveolar permeability because it is required for assembly of AJs, TJs, and the actin cytoskeleton [7,47,57–59].

MLE12 cells offer fully differentiated alveolar epithelial cells and high expansion rates for investigating these fundamental questions concerning alveolar epithelial junctions; however, lung bioengineering requires stem populations that can repopulate with more than 21 types of cells within the lungs. Several groups have identified basal epithelial stem cells (BESCs) as a promising multipotent stem cell candidate for lung regeneration [4,60] that would differentiate into ciliated, club and pneumocyte populations [61,62]. To demonstrate the potential translation of laminin treatments into stem cell repopulation of bioengineered lungs, we conducted similar coating experiments with BESCs with TEER and qPCR analysis. Like the MLE12 cells, BESCs formed higher resistance barriers on dECM and dECM with laminin compared to a complete lack of barrier formation on collagen-coated plates. Gene expression of JAM-A, EPAC, and claudin-18 were increased with both dECM and laminin compared to collagen alone. These data suggest that laminin is a stimulator of Epac and junction formation within both stem cell and fully differentiated lung epithelial cell populations.

Epac is widely known as a driver of actin organization and barrier formation within the context of endothelium, but less is known about Epac’s role in alveolar epithelial barrier formation. Initial assessment of gene expression with laminin and fibronectin showed upregulation of Epac and AF-6 with laminin coatings. Rap1 and AF6 were only upregulated in the presence of additional laminin, but not fibronectin, suggesting that laminin mediates barrier function causing upregulation of JAM-A, E-cadherin, and Claudin-18, as expected from previous literature linking Epac activation to TJ and AJ expression and stabilization [63–67]. These data led us to further investigate Epac’s role in laminin-mediated barrier function by inhibiting the interaction of Epac with downstream effectors. Epac inhibition can abolish laminin-mediated barrier resistance as observed by TEER and ZO-1 localization.

Also of interest is how we can apply Epac activation to enhance barrier formation without the addition of laminin. An Epac agonist, 007-AM was applied to collagen coatings, significantly increasing JAM-A and Rap1 gene expression, rearranging ZO-1 and the actin cytoskeleton cortically, and decreasing barrier permeability. 007-AM has recently been established as a long-lasting treatment for endothelial barriers within bioengineered lungs [7] and treatment of vascular permeability within acute lung injury [15,68]. These data now confirm that Epac activation and laminin-enriched ECM coatings can both benefit the epithelial barriers to produce functional lung biomaterials and potentially serve as therapies for edema-related diseases.

Future studies will be conducted to confirm that these findings can be translated to *ex vivo* lung tissue engineering and *in vivo* treatment of diseases. However, this high throughput and definitive *in vitro* approach has allowed us to answer basic questions about junction physiology that would not be possible within a whole lung model. Also, while these experiments have definitively established the role of both laminin and fibronectin in alveolar epithelial barrier formation, the role of other critical epithelial differentiators such as collagen IV, nidogens, GAGs, other glycoproteins will need to be investigated.

## 4. MATERIALS AND METHODS

### 4.1 Decellularization and coating preparation

Male and female porcine lungs were obtained from Smithfield-Farmland slaughterhouse or euthanized research pigs to produce dECM coating solution as previously published by our laboratory [48,69]. Quality control of dECM powders from different donors is performed by picogreen dsDNA quantification to ensure each has below 50 ng of dsDNA per mg of dry tissue. The dECM hydrogel solution is produced by pepsin digestion of 8 mg/mL dECM powder for 4 hrs. The solution was diluted to 0.1 mg/mL for use as a coating. Type I collagen from bovine skin (Sigma) diluted to 0.1 mg/mL in PBS was used as a control coating solution. The collagen and dECM solutions were also supplemented with 0.01 mg/mL laminin (Sigma, L2020) and fibronectin (Sigma, F0895) from human plasma. Collagen and dECM coating concentrations were determined by literature review and laminin and fibronectin concentrations were determined by literature review and qPCR dosing experiments (data not shown). TCP was coated overnight at 4°C to avoid gelation, then rinsed 3 times with PBS or saline (for TEER experiments) prior to cell culture.

### 4.2 Cell culture and reagents

All cells were cultured at 37 °C with 5% CO_2_. Mouse alveolar type II cell line (MLE12, ATCC) were cultured with HITES medium containing 2% fetal bovine serum according to the manufacturer’s protocols. MLE12s offer rapid doubling times and express lung surfactant proteins B and C. MLE12s were also chosen because primary differentiated human alveolar epithelial cells are not commercially available and slow proliferation of isolated cells would not be feasible. Primary human airway basal epithelial stem cells (BESCs, identified by integrin α6, KRT5, and KRT14) from healthy donors, in EpiX medium (Propagenix) were also used to demonstrate translatability to recellularization with stem cell populations [70]. Passages between 2 and 6 were used for experimentation. Longer culture periods were used for BESC experimentation to allow for differentiation and barrier formation to occur. BESCs and MLE12s on hydrogel coatings were treated with either ESI-09 (Sigma), a highly specific Epac inhibitor that does not affect PKA expression, or 8-pCPT-2-O-Me-cAMG-AM (007-AM, TOCRIS), an Epac agonist that does not discriminate between Epac1 or Epac2. 5 nM of each treatment or a DMSO vehicle control was added in media. Concentrations were determined by a dose response assay (data not shown), testing a range determined by literature review [15,71] for 8 to 24 h. Antibodies for immunofluorescent staining included, ZO-1 (Invitrogen), DAPI (Invitrogen), Alexa Fluor 488 Phalloidin (Invitrogen), and Alexa Flour 647 secondary antibody (Invitrogen).

### 4.3 Cell adhesion and spreading assay

Cell Counting Kit-8 (CCK8, Dojindo Molecular Technologies) colorimetric assay was used to determine cell viability and proliferation on each coating. 4 and 24 h after inoculating 3.2 x10^3^ MLE12 cells per cm^2^, media and unattached cells were aspirated from each well. The CCK8 and media solution was added to each well according to the manufacturer’s protocol. A microplate reader detected the absorbance of each well. Data are shown as normalized absorbance to collagen.

### 4.4 Transepithelial electrical resistance (TEER)

96W10idf PET plates (Applied Biophysics) were coated overnight at 4°C with each coating type and rinsed with media. BESCs and MLE12s were seeded at a density of 200,000 cells/cm^2^ to achieve confluence shortly after the start of culture. Cell attachment and cell-cell junction resistances were quantified using electric cell-substrate impedance sensing (ECIS, Applied Biophysics). Wells were coated and resistances were measured according to manufacturer instructions. Resistance is reported at low frequencies (4000 Hz) and longer time points to highlight cell-cell junction formation and shorter time points to show cell attachment.

### 4.5 Dextran permeability assay

Alveolar epithelial monolayers formed on 0.4 μm Transwells (Corning) coated with each ECM combination as previously stated prior to seeding of 200,000 cells/cm^2^. After 1 and 3 days, 2 mg/mL of 4 kDa FITC-dextran in PBS (Sigma) was added to the apical chamber of the Transwell for 24 h. 100 uL was taken from the bottom chamber of the well to measure fluorescent intensity using a microplate reader with excitation at 490 nm and emission at 520 nm. The intensity of media without FITC-Dextran was subtracted from each value to determine relative amounts of dextran that has passed through the barrier.

### 4.6 Gene expression quantification

mRNA was extracted and purified from MLE12s and BESCs seeded onto each coating for 5 days then treated with a DMSO vehicle control or Epac treatment for 4 h, using an RNeasy Mini kit (Qiagen) and converted to cDNA using iScript cDNA synthesis kit (Biorad). qPCR was performed with primers designed for the desired target genes shown in Table 1. After the previous examination of multiple housekeepers, 18s will be used to analyze further results.

**Table 1.**
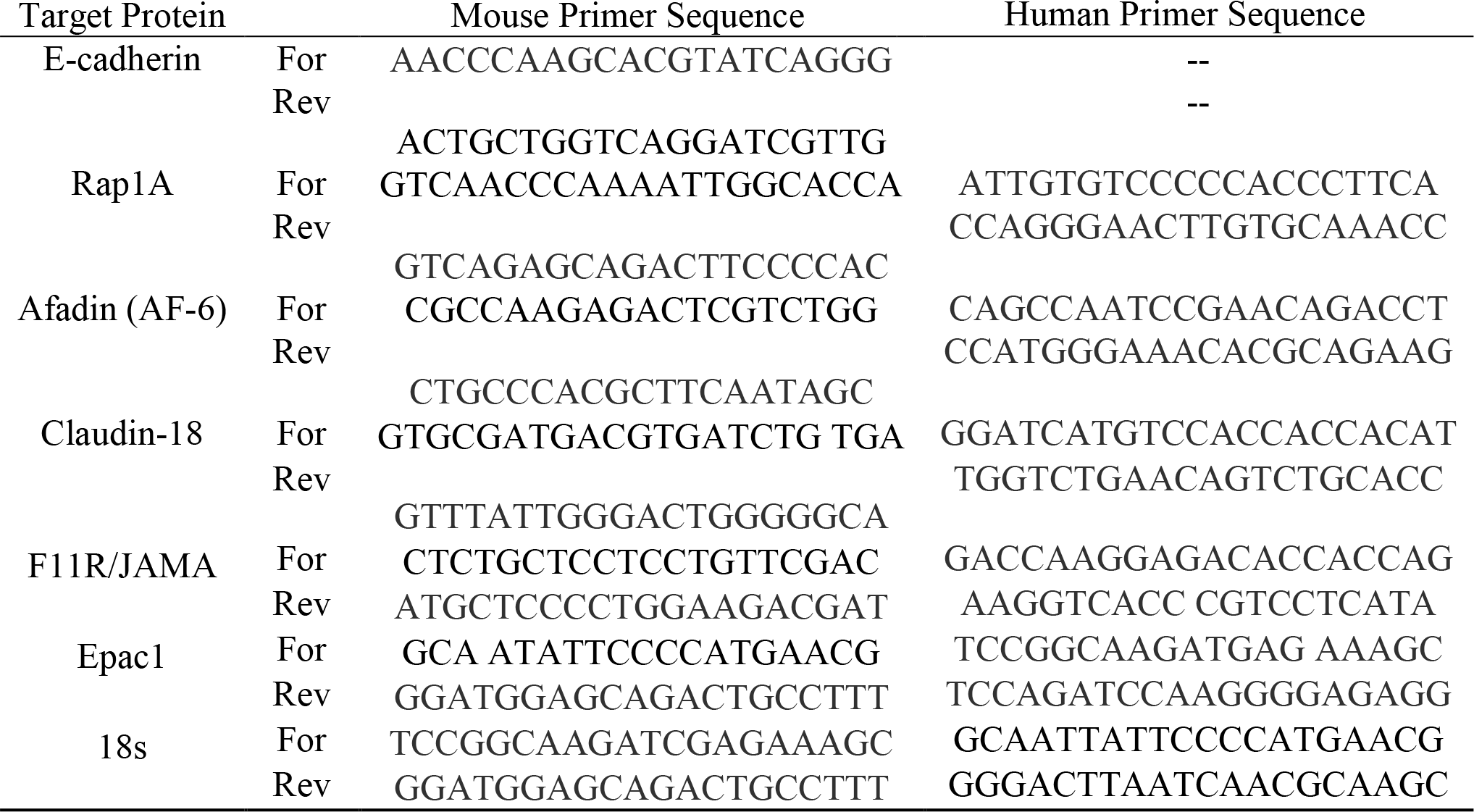
Primer sequences used in the qPCR analysis.

### 4.7 Immunofluorescence staining and ZO-1 quantification

MLE12 cells were seeded onto sterilized glass coverslips or Transwell inserts coated with each of the ECM combinations and were cultured until a confluent monolayer was achieved. Cells were treated with either ESI-09, 007-AM, or the DMSO vehicle control for 4 h. To permeabilize and fix, 0.5% Triton X-100 in 4% paraformaldehyde was applied for 2 min, then replaced with 4% paraformaldehyde for 20 min. After several rinses with PBS, the cells were blocked with 0.1% BSA before labeling with primary antibodies for 30 min at 37°C. 0.1% BSA was applied again before labeling with each with their respective secondary antibodies for 30 min at 37°C. Images will be acquired on a Zeiss LSM 710 confocal microscope using ZEN2011 software.

Quantification of the ZO-1 staining was done using the AngioTool software developed by the NIH Center for Cancer Research. A minimum of 3 images per sample and 3 samples per group were used to create a map of the junction structure and then analyze spatial parameters. The number of cells per image was counted using ImageJ software. Number of junction endpoints or breaks per cell and mean lacunarity were used to characterize junction integrity for each sample. Lacunarity quantifies the degree of spatial order within the junction structure by comparing the variation in both the background and foreground pixel densities [72,73].

## 5. SUPPLEMENTAL DATA

**Sup. Fig. 1.**
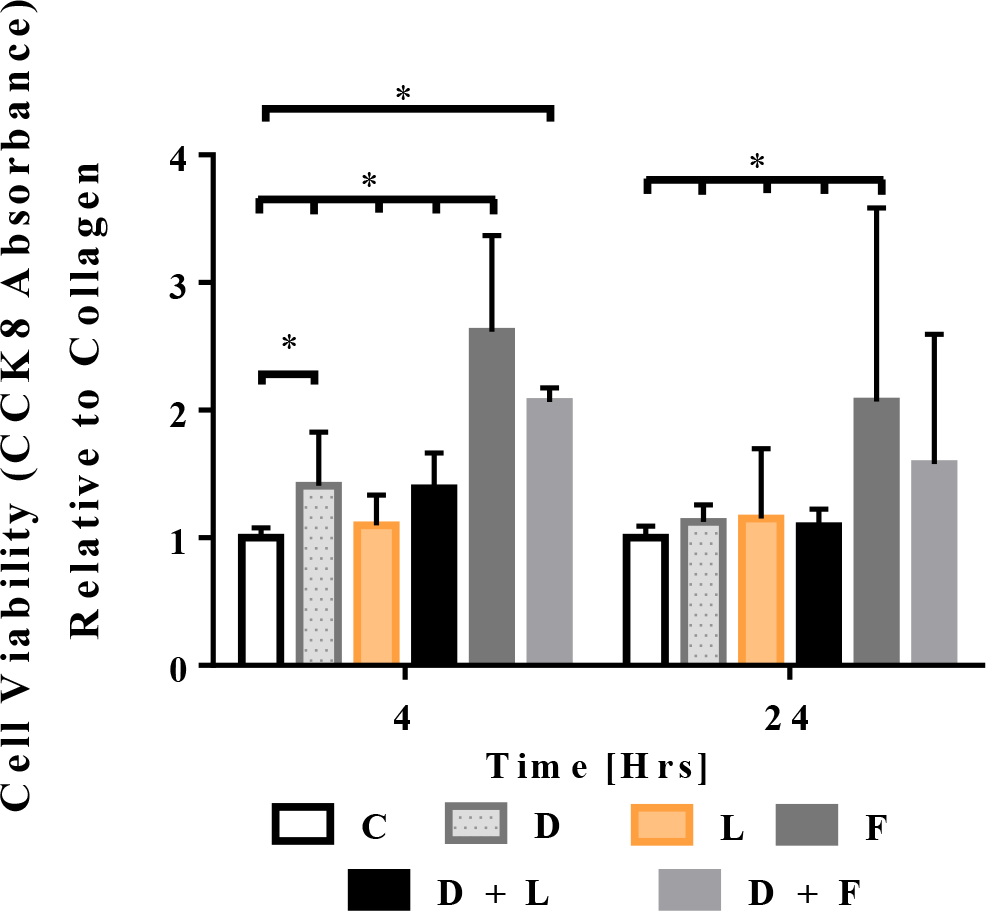
Attachment and proliferation of MLE12 cells with respect to ECM coating. Relative MLE12 cell viability on TCP coated with collagen (C), dECM (D), laminin (L), or a combination of two determined by a CCK8 assay at 4 and 24 hours. Data are presented as mean +/− standard deviation from 3 experimental replicates. *, **, *** indicates p < 0.05, p< 0.01, and p < 0.001, respectively.

## Abbreviations

ECM: extracellular matrix
dECM: decellularized extracellular matrix
TEER: trans-epithelial electrical resistance
JAM: junction adhesion molecule
ZO: zonula occludens
TJ: tight junction
AJ: adherens junction
PKA: Protein kinase A
Epac: exchange protein directly activated by cAMP
cAMP: cyclic AMP
AF-6: Afadin
EMT: Epithelial to mesenchymal transition
GAGs: glycosaminoglycans
Rap: repressor activator protein
SDC: Sodium Deoxycholate
BESC: basal epithelial stem cell
MLE12s: mouse alveolar epithelial cells

## Acknowledgments

Microscopy was performed at the VCU Microscopy Facility, supported, in part, by funding from NIH-NCI Cancer Center Support Grant P30 CA016059, and the VCU Nanomaterials Core Characterization Facility. This work was supported by the National Science Foundation (NSF CAREER CMMI 135162) and the National Institutes for Health (NIH RO1AG041823). The authors also wish to thank Nadiah Hassan for her editing and suggestions to this work.

